# Space-time home range estimates and resource selection for the Critically Endangered Philippine Eagle on Mindanao

**DOI:** 10.1101/2022.05.19.492630

**Authors:** Luke J. Sutton, Jayson C. Ibañez, Dennis I. Salvador, Rowell L. Taraya, Guiller S. Opiso, Tristan Luap P. Senarillos, Christopher J.W. McClure

## Abstract

Quantifying home range size and habitat resource selection are important elements in wildlife ecology and are useful for informing conservation action. Many home range estimators and resource selection functions are currently in use. However, both methods are fraught with analytical issues inherent within autocorrelated movement data from irregular sampling and interpretation of resource selection model parameters to inform conservation management. Here, we apply satellite telemetry and remote sensing technologies to provide first estimates of home range size and resource selection for six adult Philippine Eagles (*Pithecophaga jefferyi*), using five home range estimators and non-parametric resource selection functions. From all home range estimators, the median 95 % home range size was between 39-68 km^2^ (range: 22-161 km^2^), with the 50 % core range size between 6-13 km^2^ (range: 5-33 km^2^). The space-time autocorrelated kernel density estimate (AKDE) had the largest median 95 % home range size = 68 km^2^ and a 50 % core range = 13 km^2^. Local convex hulls (LoCoH) estimated the smallest median 95 % home range = 39 km^2^ and a 50 % core range = 6 km^2^. From the resource selection functions, all adults used areas high in photosynthetic leaf and canopy structure but avoided areas of old growth biomass and denser areas of vegetation, possibly due to foraging forays into fragmented areas away from nesting sites. For the first time, we determine two important spatial processes for this Critically Endangered raptor that can help in directing conservation management. Rather than employing a single home range estimator, we recommend that analysts consider multiple approaches to animal movement data to fully explore space-time and resource use.

## Introduction

Estimating animal home range size and habitat resource selection is a fundamental aspect in wildlife ecology and conservation (Hooten *et al*. 2017). Quantifying home range behaviour and resource selection using Global Positioning System (GPS) telemetry devices are used to inform conservation management and policy (Fieberg *et al*. 2021; Silva *et al*. 2021). Therefore, it is crucial that reliable and robust metrics are used for both. Since the inception of the home range concept (Burt 1943), many home range estimators have been used (Signer & Fieberg 2021). However, finding a reliable home range estimator has proven difficult due to the analytical challenges inherent with animal movement data that are often autocorrelated, have irregular sampling, or small sample sizes (Silva *et al*. 2021). Similarly, estimating resource selection functions by comparing environmental covariates at an individual’s used locations to those environmental locations assumed to be available with logistic regression is popular (Johnson *et al*. 2006). However, interpreting resource selection model parameters to inform management is difficult (Fieberg *et al*. 2021).

An animal’s home range is formally defined as those movements regularly used for foraging and breeding but excluding occasional sallies outside of this area (Burt 1943; Fieberg & Borger 2012). Thus, an animal’s home range reflects its ecological needs and the decisions that result from these environmental requirements (Tétreault & Franke 2017). Home ranges are therefore expected to differ amongst individuals within a species over space and time dependent on shifting ecological needs and varying resources (Signer & Fieberg 2021). Further, selection of a specific home range estimator can in itself explain as much of the variation in home range size as the ecological processes influencing it (Signer *et al*. 2015; Tétreault & Franke 2017).

Home range estimators can be split into two classes: geometric and probabilistic (Signer & Fieberg 2021). Geometric estimators are built following a set of hull-based rules, with a typical example being a minimum convex polygon (MCP). However, the MCP often overestimates the home range, with Local Convex Hulls (LoCOH, Getz & Wilmers 2004), which generalize the MCP, an improved estimator able to account for autocorrelation, better reflecting the true home range by considering hard boundaries within the range extent (Getz *et al*. 2007). Further, Time Local Convex Hulls (T-LoCoH) are a further generalization of local convex hulls, incorporating time by using adaptive scaling of individual velocities to define a utilization distribution that captures space-time use (Lyons *et al*. 2013). Conversely, probabilistic estimators are constructed using an underlying probabilistic model which estimates a utilization distribution, that is, the relative frequency distribution of an animal’s locations in two-dimensional space (Van Winkle 1975). The utilization distribution is an extension of the original home range concept (Burt 1943), where an animal’s use of space is defined by a probability density function that quantifies the chance the animal will be found at any given location within its home range (Van Winkle 1975; Worton 1987).

Kernel density estimators (KDEs, Worton 1989) are a non-parametric probabilistic estimator, fitted with both fixed and adaptive kernel bandwidths to account for over smoothing (Wand & Jones 1994). However, fixed and adaptive KDEs can overestimate home range sizes, even when accounting for bandwidth over smoothing with an adaptive kernel (Silva *et al*. 2021). Recently, autocorrelated kernel density estimates (AKDE, Fleming & Calabrese 2017) have been proposed as an improvement on fixed and adaptive KDEs. AKDEs first fit an Ornstein-Uhlenbeck (Uhlenbeck & Ornstein 1930) continuous-time stochastic process movement model to the animal locations, and then incorporate the movement model into an area-corrected home range estimator with weighting that accounts for autocorrelation and irregular sampling (Calabrese *et al*. 2016; Silva *et al*. 2021). Space-time home range estimates are therefore expected to provide more robust estimates of the utilization distribution because they account for the important third dimension of time in animal movement patterns (Keating & Cherry 2009).

Within an animal’s home range, resource selection functions (RSFs) are used to infer the probability of resource use for a given individual within that defined area (Manly *et al*. 2002). Standard parametric logistic regression is the most popular method to quantify resource selection (Johnson *et al*. 2006) but has been criticized because used locations (species presence) are continuous points but are compared to available locations (raster pixels) in discrete space (Keating & Cherry 2004; Fieberg *et al*. 2021). Poisson point processes have been proposed as an alternative to standard parametric resource selection functions to make habitat selection analyses easier to understand and more accessible to a wide range of end users (Baddeley *et al*. 2012). For ease of interpretation, non-parametric RSFs can be fitted directly to the species locations without accounting for available locations using a point process intensity probability density function based on a kernel density estimate (Baddeley *et al*. 2012).

The Philippine Eagle (*Pithecophaga jefferyi*) is a globally threatened tropical forest raptor (Bildstein *et al*. 1998), currently classified as ‘Critically Endangered’ on the IUCN Red List (BirdLife International 2018). This large eagle is endemic to four islands in the Philippine archipelago (Mindanao, Leyte, Samar, and Luzon; Kennedy 1977), with a restricted distribution across lowland and montane tropical forests (Salvador & Ibañez 2006; Sutton *et al*. 2022). The two key threats to its future survival are habitat loss and human persecution (Salvador & Ibañez 2006). Despite its elevated extinction risk, fundamental aspects of Philippine Eagle ecology such as home range size and habitat use are relatively unknown. Indeed, the IUCN Red List suggests that further research into ecological requirements is urgently required to inform conservation actions (BirdLife International 2018). Here, we use satellite telemetry locations from six GPS tagged adult Philippine Eagles to (**1**), estimate home range size using five geometric and probabilistic estimators, and (**2**), quantify habitat use with non-parametric resource selection functions. Finally, we outline how quantifying these key ecological processes can inform conservation action for this raptor of conservation concern.

## Methods

### GPS telemetry data

We sourced Philippine Eagle satellite telemetry locations from the Philippine Eagle Foundation that is archived in the Global Raptor Impact Network (GRIN, McClure *et al*. 2021), a data information system for global population monitoring for all raptors. For the Philippine Eagle, GRIN includes GPS fixes from six breeding adult Philippine Eagles (four females, two males) on the island of Mindanao. All Philippine Eagles were trapped using either a modified Bal-Chatri (Miranda & Ibanez 2006) or a large bownet baited with domestic rabbit (*Oryctolagus cuniculus*). Two eagles were instrumented with solar-powered Global Positioning System-Global System for Mobile Communications (GPS-GSM) transmitters (weight = 70 g; Microwave Telemetry, Inc) while four eagles had battery-powered LC4™ Argos-GPS platform transmitter terminal (PTT) fitted (weight = 105g; Microwave Telemetry, Inc), harnessed with Teflon-coated nylon ribbon backpacks. All tags weighed < 3 % of the body weight for all adults tagged. Tags were programmed to transmit on a 2-hr sampling interval for adults 001F, 002F, 004M, 006F, with adult 003F at 24 hrs and adult 005M at 2 mins. All birds were marked with aluminium leg bands – the four females with blue bands on their left tarsus, and the two males with green bands on their right tarsus. All GPS transmitter harnessing was conducted with a Gratuitous Permit to trap and tag the birds in the presence of a veterinarian as required by the national government of the Philippines.

A total of 80,481 fixes were obtained from four adult females and two adult males from April 2013 to September 2021 (Table 1). We removed all duplicated records and used all raw GPS fixes in the autocorrelated kernel density estimates (AKDEs) for all birds expect 005M which we sub-sampled using a 3-hr interval due to computing constraints using the full raw dataset of 74,098 fixes. For the fixed and adaptive kernel density estimates (KDEs) along with local convex hull (LoCoH) estimators, we subsampled fixes from all birds using a minimum 3-hour interval between fixes to achieve consistency across estimators and to account for autocorrelation (Signer & Fieberg 2021). We assessed how effective the number of GPS relocations was at capturing the utilization distribution using an incremental analysis with bootstrapped minimum convex polygons (*n =* 100), quantifying when the number of relocations within the MCP area reached an asymptote (Walls & Kenward 2012), using the ‘hrBootstrap’ function in the R package move (Kranstauber *et al*. 2020). From our bootstrapped estimates, the number of relocations for all six adults was sufficient at capturing the MCP utilization distribution, ranging from asymptotes of 100 relocations for adult 003F to 1000 relocations for adult 005M (Fig. S1).

**Table 1.**
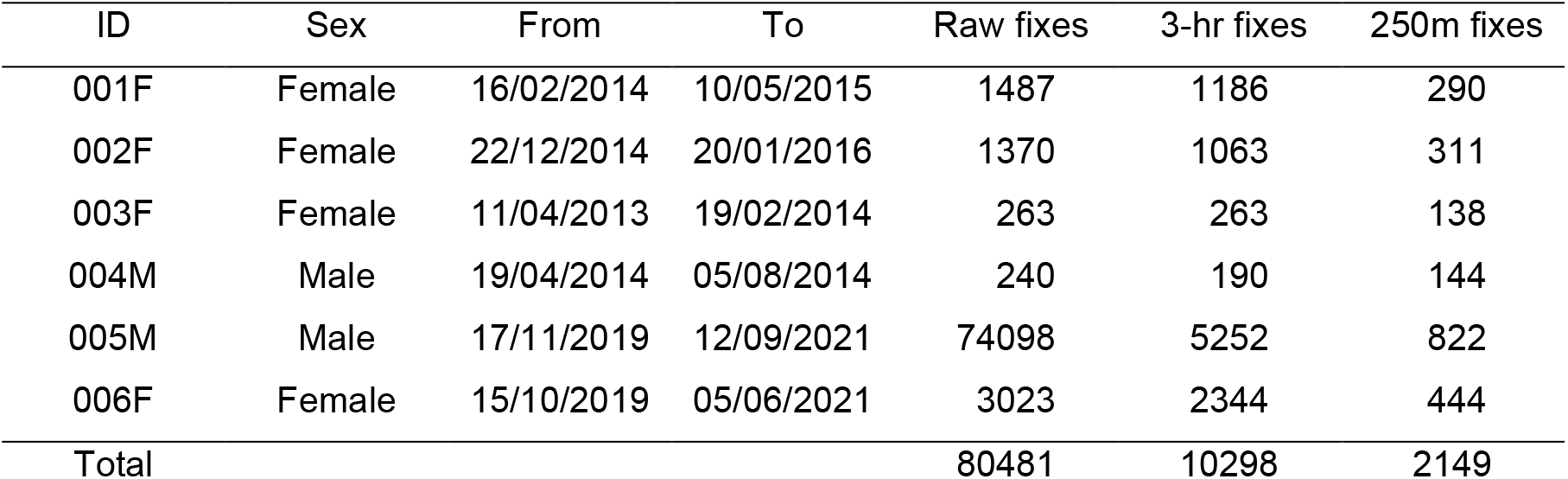
Global Positioning System (GPS) telemetry metadata for six satellite tagged adult Philippine Eagles from the island of Mindanao, used for home range estimation. Totals for 3-hr fixes are subsampled from the raw data locations using a 3-hr sampling rate interval. Totals for 250m fixes are the number of spatially thinned fixes using a 250m spatial filter.

| ID | Sex | From | To | Raw fixes | 3-hr fixes | 250m fixes |
| --- | --- | --- | --- | --- | --- | --- |
| 001F | Female | 16/02/2014 | 10/05/2015 | 1487 | 1186 | 290 |
| 002F | Female | 22/12/2014 | 20/01/2016 | 1370 | 1063 | 311 |
| 003F | Female | 11/04/2013 | 19/02/2014 | 263 | 263 | 138 |
| 004M | Male | 19/04/2014 | 05/08/2014 | 240 | 190 | 144 |
| 005M | Male | 17/11/2019 | 12/09/2021 | 74098 | 5252 | 822 |
| 006F | Female | 15/10/2019 | 05/06/2021 | 3023 | 2344 | 444 |
| Total |  |  |  | 80481 | 10298 | 2149 |

To test for range residency we calculated semi-variance functions visualised with empirical variograms to identify unbiased estimates of stationary movement periods of site fidelity with data containing time-averaged autocorrelation structure in the R package ctmm (Calabrese *et al*. 2016). Variograms represent the average square distance travelled within a specified time lag. We used a median sampling interval for the time lag bin widths and Markovian Confidence Intervals for calculating the maximum number of non-overlapping lags (Calabrese *et al*. 2016). All six adults showed site fidelity with clear asymptotes ranging between 2 to 18 km continuous range residency behaviour after 3 to 9 day short time lags and all less than one calendar month from tagging (except adult 006F which was less than 2 calendar months), supporting the application of home range analysis (Figs. S2-S3).

### Home range estimation

Utilization distributions were constructed to estimate the probability of relocating an individual within a given home range using the standard definitions in two-dimensional space (Van Winkle 1975; Worton 1987, 1989), which we further extend to three-dimensional space-time (Keating & Cherry 2009). We calculated utilization distributions using five home range estimators because of variation in outputs between different estimator methods (Signer & Fieberg 2021). For all estimators we fitted 95 % probability of use contour isopleths to represent the home range utilization distribution (Laver & Kelly 2008), and 50 % probability of use contour isopleths to represent a core range utilization distribution, characteristic of a territorial area (White & Garrott 1990). We selected a core range of 50 % probability of use to maintain consistency across the different estimators but recognise that defining a 50 % core range is not always appropriate (Vander Wal & Rodgers 2012). All home range area estimates were calculated in a Universal Time Mercator (UTM) projection in R (v3.5.1; R Core Team 2018) and following analytical recommendations from Laver & Kelly (2008).

### Kernel Density Estimates

We calculated utilization distributions using three different kernel density estimators (Worton 1989). First, we fitted standard fixed bandwidth non-normal Epanechnikov kernels (Epanechnikov 1969), with an ad-hoc reference smoothing parameter (*h*_*ref*_) multiplied by 1.77 (Silverman 1986), based on the number of locations and the variance between x and y coordinates. Second, we fitted adaptive smoothing plug-in bandwidth bivariate kernel estimates (*h*_*pi*_) (Wand & Jones 1994) using a Sum of the Asymptotic Mean Squared Error (SAMSE) pilot bandwidth selector (Duong & Hazelton 2003). We assessed a range of potential univariate plug-in bandwidth selectors (termed ‘pilots’) and opted for SAMSE due to its higher numerical stability (Duong 2007) and the low variance between each respective pilot bandwidth. We fitted both fixed and adaptive KDEs in the R packages adehabitatHR (Calenge 2006), ks (Duong 2007) and sp (Bivand *et al*. 2013), with R code adapted from Tétreault & Franke (2017).

We fitted autocorrelated KDEs (AKDEs; Fleming & Calabrese 2017) in the R package ctmm (Calabrese *et al*. 2016) with a movement model that best explains the autocorrelated structure of our tracking data using a perturbative Hybrid Residual Maximum Likelihood parameter estimator (pHREML), which is a form of maximum likelihood estimation that reduces bias in variance/covariance estimation (Silva *et al*. 2021). AKDEs were fitted with a continuous-time stochastic process movement model to overcome the autocorrelated nature of our GPS tracking fixes and mitigate small absolute and effective sample sizes (Calabrese *et al*. 2016). We evaluated a pool of candidate movement models for each individual eagle from Ornstein-Uhlenbeck movement patterns including both isotropic (symmetrical diffusion) and anisotropic (asymmetrical diffusion) variants, along with the standard KDE assumption of independent and identical distributed (IID) data, based on Akaike’s Information Criterion (Akaike 1974) adjusted for small sample sizes (AICc; Hurvich & Tsai 1989). We considered all models with a ΔAICc < 2 as having strong support (Burnham & Anderson 2004). From our candidate models, the best supported movement process for all six eagles was an Ornstein-Uhlenbeck anisotropic process (ΔAICc = 0.0; Table S1), which we then fitted into an area-corrected AKDE home range estimator with additional weighting that upweights fixes in under-sampled times and down-weights fixes in over-sampled times (Silva *et al*. 2021).

### Local Convex Hulls

We calculated utilization distributions using fixed and temporal Local Convex Hull (LoCoH) estimators, both using *k* nearest neighbour convex hulls, which are a generalization of a minimum convex polygon estimator (Getz & Wilmers 2004), in the R package tlocoh (Lyons *et al*. 2013). We constructed fixed local convex hulls by associating each point and its *k* -1 nearest neighbours localized in space. The hulls were then ordered smallest to largest, taking the cumulative union of each hull from smallest upwards thus constructing the utilization distribution isopleths, with the smallest hulls indicating the most frequently areas, i.e., the 10% isopleth contains 10% of the points with a higher utilization than the 95% isopleth that contains 95% of the points (Getz *et al*. 2007; Tétreault & Franke 2017). In addition, we constructed time local convex hulls (T-LoCoH), which are a generalization of LoCoH, incorporating time into the aggregation of the *k*-nearest neighbour local convex hulls in Euclidean space using adaptive scaling of individual velocities to define a utilization distribution that captures space-time use (Lyons *et al*. 2013). T-LoCoH incorporates the timestamp as a time-scaled distance metric between any two points into a third axis of Euclidean space in the selection of *k*-nearest neighbours and hull sorting within the LoCoH algorithm.

### Resource Selection

#### Habitat covariates

We quantified resource selection using the GPS fixes and three habitat covariates derived from satellite remote sensing data using 16-day 250-m composite surface reflectance band imagery from the Moderate Resolution Imaging Spectroradiometer (MODIS, https://modis.gsfc.nasa.gov/) product MCD13Q1. We used two surface reflectance bands that represent unclassified raw measures of vegetation structure and composition, used previously to represent vegetation patterns (Morán-Ordóñez *et al*. 2012; Shirley *et al*. 2013; Van doninck *et al*. 2020). Band 2 Near Infrared represents leaf and canopy structure, with Band 7 Short Wave Infrared related to senescent or old growth biomass (Shirley *et al*. 2013). Additionally, we used Enhanced Vegetation Index (EVI) processed using all four MODIS surface reflectance bands using the ‘spectralIndices’ function in the R package RStoolbox (Leutner *et al*. 2019). EVI ranges on a scale from -1 to 1, with positive values closer to 1 indicating dense, healthy vegetation, and negative values indicating low vegetation cover.

EVI is an optimized vegetation index responsive to canopy structure variations and with improved sensitivity in areas of high biomass through reduction in background noise and atmospheric influences (Huete *et al*. 2002). We selected EVI due to its superior performance at capturing dense vegetation characteristics and canopy structure in tropical regions compared to other spectral indices such as Normalised Difference Vegetation Index (NDVI; Qiu *et al*. 2018), which tends to saturate in densely vegetated areas (Huete *et al*. 2002). We downloaded imagery corresponding to the start and end dates over the time period of each tracked eagle using the R package MODIStsp (Busetto & Ranghetti 2016) and calculated mean surface reflectance values over each respective time period to use in processing the covariates. All surface reflectance bands contain spectral reflectance values estimated by target at surface, calibrated with cloud detection and atmospheric corrections. Reflectance values are expressed as the ratio of reflected over incoming radiation, meaning reflectance can be measured between the values of zero and one. Absolute reflectance values of 3-4 indicate healthy vegetation (Huete *et al*. 2004). All covariates used for each respective eagle had low collinearity with Variance Inflation Factors <2.

#### Resource Selection Functions

We thinned GPS fixes using a 250-m spatial filter (Table 1) to match the resolution of the covariate rasters and fitted presence points and the three covariates to individual RSFs following third-order home range resource selection (Johnson 1980). We defined a resource use home range for each individual eagle by merging the 95 % maximum likelihood AKDE with a 100 % minimum convex polygon to fully capture the total potential home range and thus all the spatially filtered GPS fixes therein (Northrup *et al*. 2013). We fitted non-parametric RSFs where we only considered resource use at presences using a point process intensity probability density function using the ‘rhohat’ function in the R package spatstat (Baddeley & Turner 2005). RSFs were fitted by computing a non-parametric kernel smoothing estimate of locations as a point process intensity function λ (*u*) of the three spatial covariates over each respective eagles’ home range window following the formulation of Baddeley *et al*. (2012),

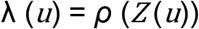

where *Z* is the spatial covariate and *ρ* (*z*) is the resource selection function to be estimated, with *u* representing location. We fitted Gaussian kernel densities with variable-bandwidth kernel smoothing using cross-validated bandwidth selection which assumes a Cox process for clustered data (Diggle 1985) and an isotropic edge correction for polygon windows derived from Ripley’s K-function (Ripley 1988). Additionally, we corrected for sampling bias with Horvitz-Thompson weighting (Horvitz & Thompson 1952), where each GPS fix in the sample is weighted by the reciprocal of its sampling probability. We fitted all RSFs with 95 % Confidence Intervals.

## Results

### Home Range Estimation

#### Kernel Density Estimates

The median 95 % home range estimate from the fixed Epanechnikov KDE was 61 km^2^ (SE ±13.5), and the median 50 % core home range estimate 12 km^2^ (SE ±1.9), with the core range comprising 21 % of the 95% home range area (Table 2; Fig. S4). Home range estimates from the adaptive SAMSE KDE were smaller, with the median 95 % home range estimate 43 km^2^ (SE ±5.7) and a median 50 % core home range estimate of 7 km^2^ (SE ±1.2), with the core range comprising 19 % of the 95% home range area (Table 2; Fig. S4). The median 95 % home range estimate from the weighted AKDEs was greater than both the fixed and adaptive estimates at 68 km^2^ (CI = 62-74 km^2^), with the median 50 % core home range estimate 13 km^2^ (CI = 11-14 km^2^), comprising 21 % of the 95% home range area (Table 3, Fig. 1).

**Table 2.** Fixed and adaptive kernel density home range estimates (KDE) for six adult Philippine Eagles on the island of Mindanao. Estimates calculate 95 % probability of use contour isopleths to represent the home range utilization distribution and 50 % probability of use contour isopleths to represent a core range utilization distribution, SAMSE = Sum of the Asymptotic Mean Squared Error pilot bandwidth selector. All area values in the 95% and 50% columns are km^2^.

| ID | Epanechnikov fixed KDE |  |  | SAMSE adaptive plug-in KDE |  |  |
| --- | --- | --- | --- | --- | --- | --- |
|  | 95% | 50% | % core | 95% | 50% | % core |
| 001F | 58 | 11 | 19 | 37 | 7 | 19 |
| 002F | 64 | 13 | 20 | 48 | 9 | 19 |
| 003F | 43 | 9 | 21 | 28 | 7 | 25 |
| 004M | 87 | 20 | 23 | 58 | 13 | 22 |
| 005M | 36 | 8 | 22 | 26 | 5 | 19 |
| 006F | 126 | 17 | 13 | 56 | 6 | 11 |
| Median | 61 | 12 | 21 | 43 | 7 | 19 |

**Table 3.** Autocorrelated kernel density estimates (AKDE) for six adult Philippine Eagles on the island of Mindanao. Estimates calculate 95 % probability of use contour isopleths to represent the home range utilization distribution and 50 % probability of use contour isopleths to represent a core range utilization distribution with 95% Confidence Intervals (CI). All area values in the 95% and 50% columns are km^2^.

| ID | Autocorrelated KDE |  |  |
| --- | --- | --- | --- |
|  | 95% (CI) | 50% (CI) | % core |
| 001F | 64 (59-70) | 12 (11-13) | 19 |
| 002F | 71 (64-78) | 13 (12-14) | 18 |
| 003F | 39 (33-45) | 9 (8-11) | 24 |
| 004M | 108 (85-133) | 24 (18-29) | 22 |
| 005M | 41 (37-46) | 9 (8-10) | 22 |
| 006F | 161 (133-192) | 33 (28-40) | 21 |
| Median | 68 (62-74) | 13 (11-14) | 21 |

**Figure 1.**
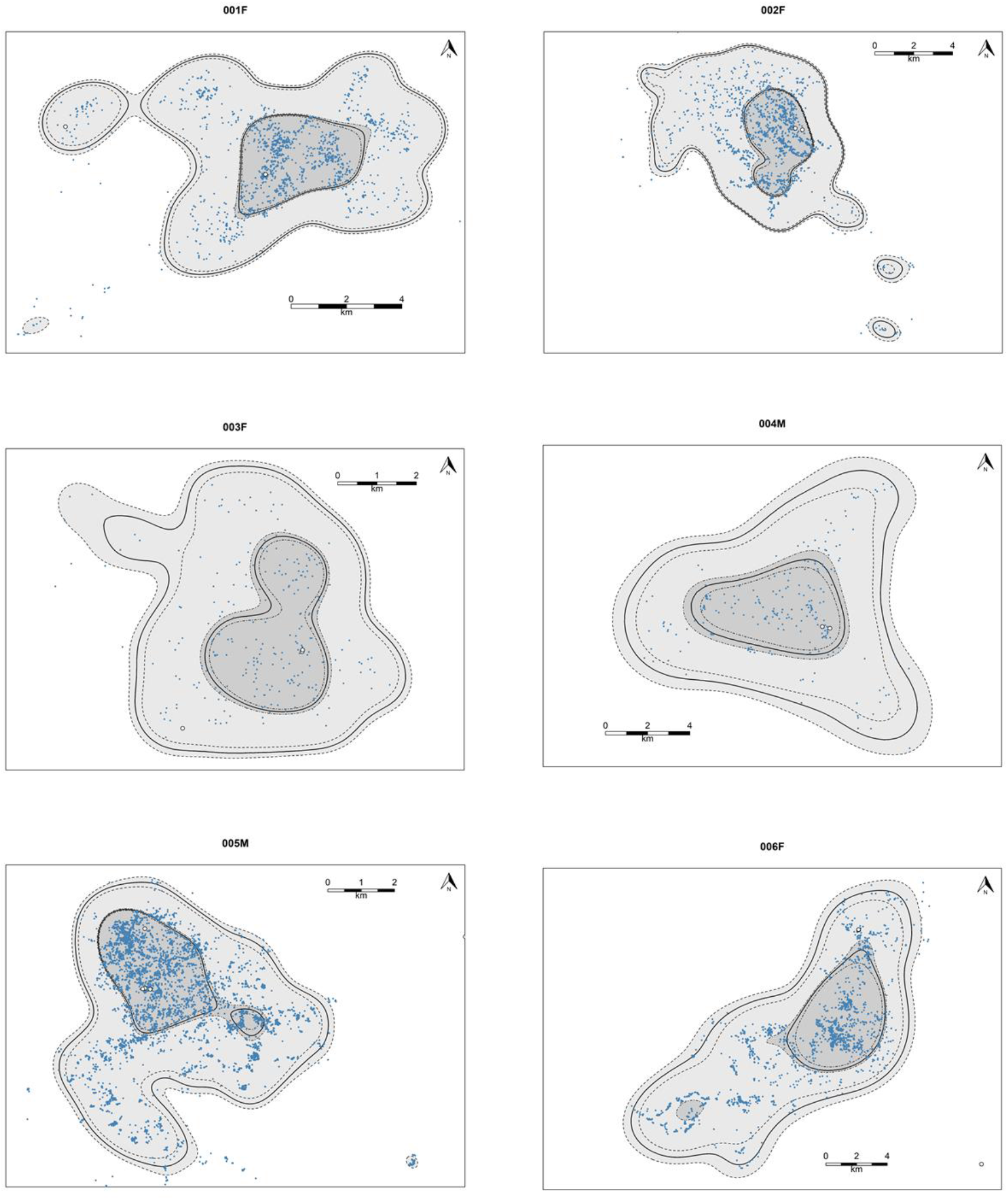
Autocorrelated kernel density estimates (AKDE) for six adult Philippine Eagles on the island of Mindanao. Maximum likelihood estimates (bold black lines) calculate 95 % probability of use (light grey) to represent the home range utilization distribution and 50 % probability of use (dark grey) to represent a core range utilization distribution. Hashed lines show 95% Confidence Intervals for both home and core range maximum likelihood estimates. Blue points show raw locations for each respective adult Philippine Eagle, except for adult 005M which was sub-sampled using a 3-hr interval due to computing constraints. White points indicate nest sites.

#### Local Convex Hulls

The median 95 % home range estimate from the LoCoH estimators was 39 km^2^ (SE ±7.8), and the median 50 % core home range estimate 6 km^2^ (SE ±0.8), comprising 20 % within the 95% home range area (Table 4; Fig. S5). Home range estimates from the T-LoCoH were larger, with the median 95 % home range estimate 56 km^2^ (SE ±12.0) and the median 50 % core home range estimate 13 km^2^ (SE ±1.2), comprising 25 % of the 95% home range area (Table 4; Fig. 2). Overall, using the median estimates there was a 19-21 % probability of space use within the 50 % core range across all estimators, except for the temporal LoCoH where 50 % probability of use increased to 25 % core range use (Table 4). AKDE estimated the largest range of 95 % utilization distributions (39-161 km^2^), with the adaptive KDE estimating the smallest range of 95 % utilization distributions (26-58 km^2^, Fig. 3). Adult female 003F and adult male 005M had the narrowest range of home range size estimates (Fig. 3), with adult female 006F having the broadest range of home range size estimates overall (Figs. 3 & 4).

**Table 4.** Local Convex Hull (LoCoH) and time Local Convex Hull (T-LoCoH) home range estimates for six adult Philippine Eagles on the island of Mindanao. Estimates calculate 95 % probability of use contour isopleths to represent the home range utilization distribution and 50 % probability of use contour isopleths to represent a core range utilization distribution. All area values in the 95% and 50% columns are km^2^.

| ID | LoCoH |  |  | T-LoCoH |  |  |
| --- | --- | --- | --- | --- | --- | --- |
|  | 95% | 50% | % core | 95% | 50% | % core |
| 001F | 33 | 8 | 24 | 60 | 13 | 22 |
| 002F | 49 | 10 | 20 | 53 | 14 | 26 |
| 003F | 22 | 5 | 23 | 24 | 9 | 38 |
| 004M | 45 | 7 | 16 | 58 | 16 | 28 |
| 005M | 26 | 5 | 19 | 34 | 8 | 24 |
| 006F | 74 | 5 | 7 | 109 | 13 | 12 |
| Median | 39 | 6 | 20 | 56 | 13 | 25 |

**Figure 2.**
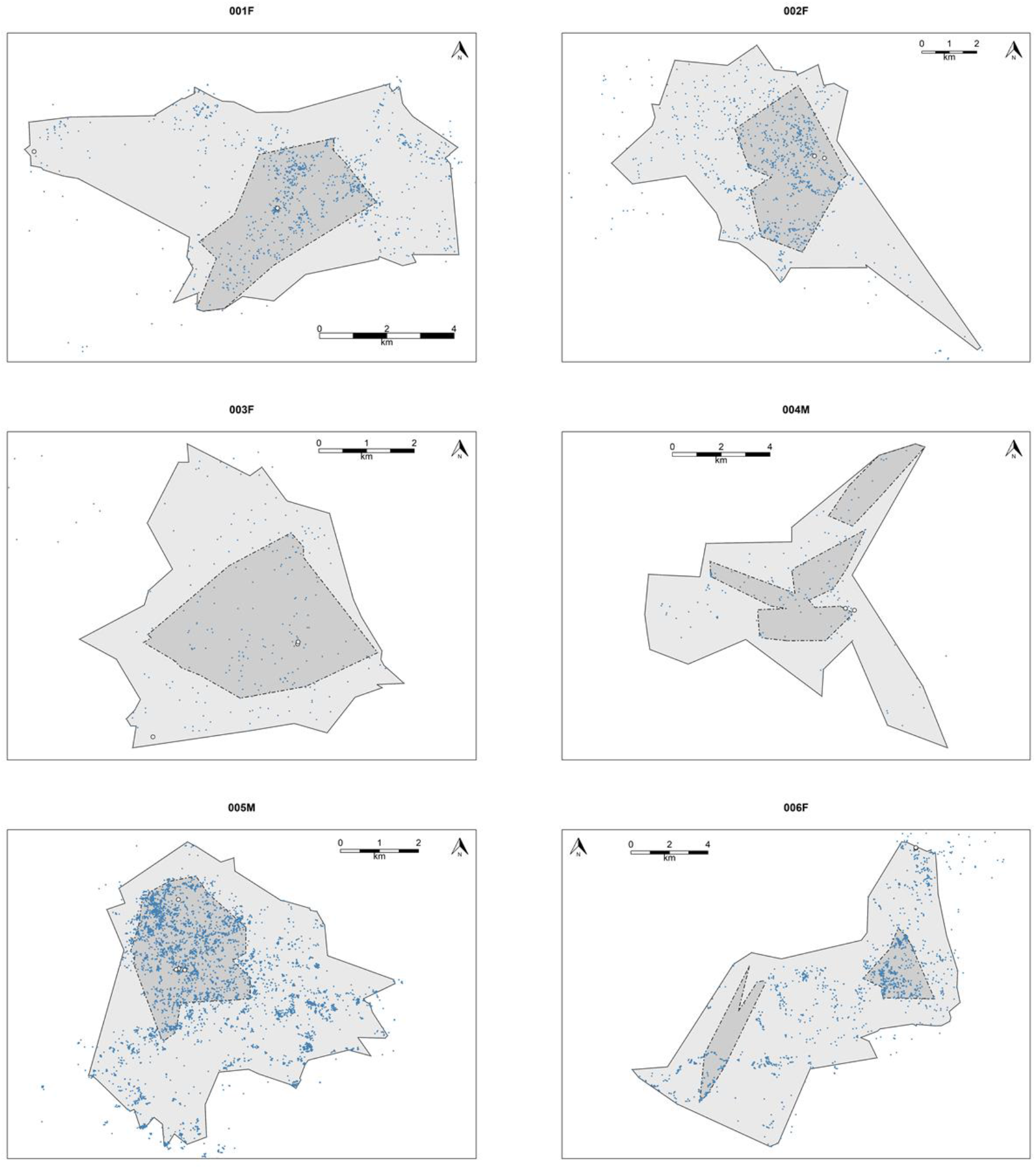
Time Local Convex Hull (T-LoCoH) home range estimates for six adult Philippine Eagles on the island of Mindanao. Estimates calculate 95 % probability of use to represent the home range utilization distribution (light grey) and 50 % probability of use to represent a core range utilization distribution (dark grey). Blue points show filtered locations using a 3-hr sampling interval for each respective adult Philippine Eagle. White points indicate nest sites.

**Figure 3.**
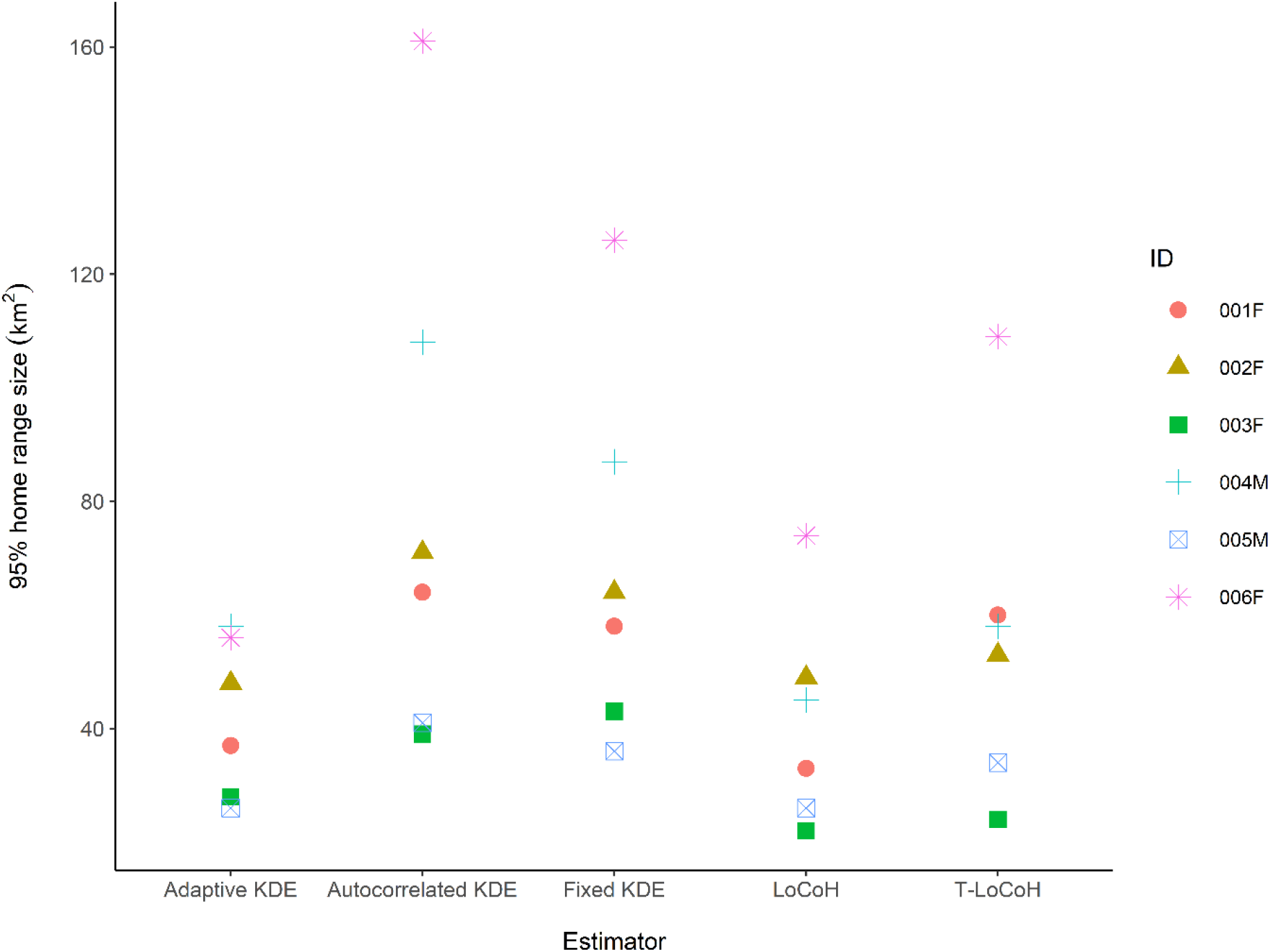
Comparison of five home range size estimators from 95 % probability of use to represent the home range utilization distribution for six adult Philippine Eagles on the island of Mindanao. KDE = kernel density estimate, LoCoH = Local Convex Hull, T-LoCoH = Time Local Convex Hull.

**Figure 4.**
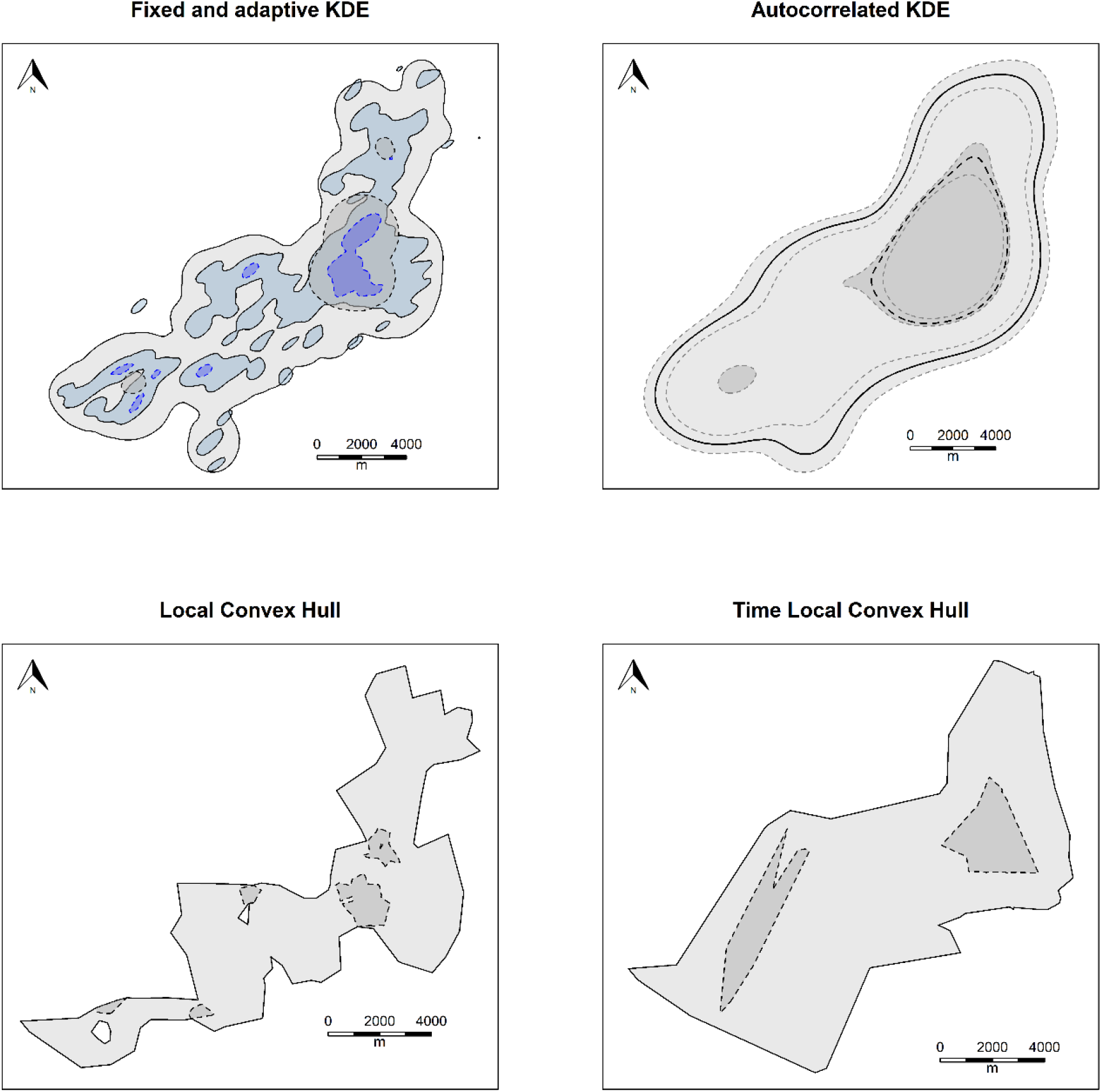
Home range estimates for adult female Philippine Eagle 006F using five home range estimators (KDE = kernel density estimate). Estimates calculate 95 % probability of use to represent the home range utilization distribution (light grey with solid black lines) and 50 % probability of use to represent a core range utilization distribution (dark grey with hashed black lines), except for the adaptive KDE 95 % home range which is shown in light blue with solid black line and 50 % core range shown in dark blue with hashed black line. For the autocorrelated KDE 95 % Confidence Intervals are shown by hashed light grey lines.

### Resource selection

From the non-parametric RSF response functions, all six eagles were associated with Band 2 Near Infrared values peaking between 0.34-0.39 (Fig. 5), indicating a relationship with dense, healthy leaf and canopy structure. Band 7 Shortwave Infrared values peaked between 0.07-0.14, indicating an association with areas of lower percent old growth biomass for all adults (Fig. 5). All six adults were more likely to be associated with EVI values between 0.35-0.55 (Fig. 5), indicating resource use of moderately dense vegetation averaged over the annual vegetation growth cycle.

**Figure 5.**
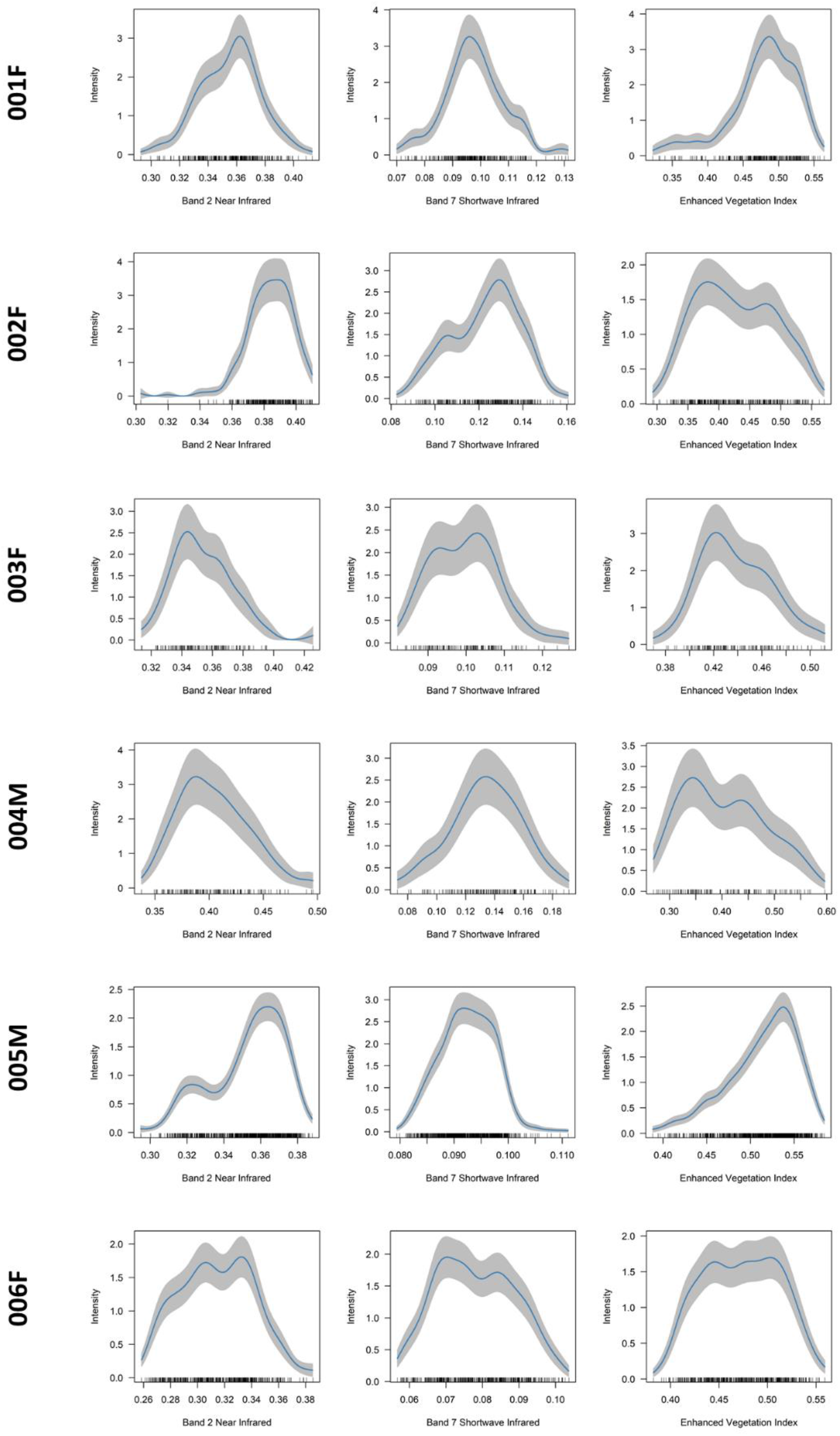
Non-parametric resource selection response curves (blue lines) using point process intensity probability density functions for six adult Philippine Eagles on the island of Mindanao. Grey shading represents 95% Confidence Intervals.

## Discussion

Quantifying animal space and habitat use is fundamentally important in understanding the ecological processes influencing an individual animal’s behaviour and movement (Hooten *et al*. 2017). By using a suite of home range estimators, our results demonstrate that adult Philippine Eagles on Mindanao have relatively small home ranges, with 75-80 % of space-time use outside of their core territorial range. AKDE estimated the largest median 95 % home range size = 68 km^2^ and the largest median 50 % core range = 13 km^2^. LoCoH estimated the smallest median 95 % home range = 39 km^2^ and the smallest 50 % core range = 6 km^2^. Additionally, most adults used areas high in photosynthetic leaf and canopy structure but tended to avoid areas of old growth biomass and denser areas of vegetation, possibly due to extended foraging movements outside of densely forested nesting areas. Our results quantify for the first time two key ecological processes for this critically endangered raptor that can be useful in informing conservation management.

### Home Range Estimation

Although the median home range estimates for all adults combined was between 39-68 km^2^ for the 95 % home range and 6-13 km^2^ for the 50 % core range, there was wide variance in home range sizes for each individual eagle irrespective of the estimator used (see Fig. 3). For example, variance amongst the adaptive 95 % kernel estimates was lower (range = 28-56 km^2^), compared to the fixed 95 % kernels (range = 36-126 km^2^), with the 95 % LoCoH hulls having lower variance (range = 22-74 km^2^), compared to the T-LoCoH hulls (range = 24-109 km^2^). Though we did not test this directly, we assume that high variance in home range size amongst individual eagles is driven by varying resource needs for each eagle across fragmented forest on Mindanao. The ratio of percent space use for the 50 % core range within the 95 % home range was generally consistent across all estimators between 19-21 %, except for T-LoCoH where this increased to 25 % core range use. Thus, adult Philippine Eagles are using 75-80 % of space-time use outside of the core territorial area, presumably when searching for food within their home range.

Previous home range estimates for the Philippine Eagle calculated median 95 % home range sizes between 64-90 km^2^ (Sutton *et al*. 2022), similar to our estimates here. These uniform estimates are not surprising because Sutton *et al*. used the same satellite telemetry dataset to calculate home range sizes but using a fixed Gaussian KDE, a radius LoCoH and a minimum convex polygon as estimators. Prior to these quantitative home range estimates, Rabor (1968) suggested a home range of 40-50 km^2^ for the Philippine Eagle, within the lower range of our median 95 % estimates, with Gonzales (1968, 1971) suggesting up to 100 km^2^. However, Kennedy (1977) calculated much lower home range sizes of between 13-25 km^2^ based on polygon and circular estimates from observer sightings of a pair of breeding eagles within an approximately 5×5 km^2^ area. Assuming these sightings were of a nesting territorial pair then they are remarkably similar to our upper range of 50 % core territorial range estimates.

### Resource selection

Habitat resource selection by animals will often give contrasting results related to issues of scale (Boyce 2006). Our results showed all eagles were associated with medium Band 2 Near Infrared reflectance values, representing healthy photosynthetic leaf and canopy structure but low Band 7 Shortwave Infrared values representing old growth forest, in contrast to a previous range-wide habitat use assessment (Sutton *et al*. 2022). Thus, solely using GPS fixes from the six adults captured the finer scale home range resource use, which is generally outside of old growth forest areas. This is possibly related to adults foraging over secondary forest and cleared agricultural lands along forest edges (Kennedy 1977; Salvador & Ibañez 2006). These foraging areas are distant from nest sites which are generally within denser forested areas (Salvador & Ibañez 2006; Ibañez *et al*. 2003). This assumption is further supported by the general association with medium values of Enhanced Vegetation Index, indicating most adults are using areas of canopy vegetation density between EVI values of 0.35-0.55 over the annual growth period (see Fig. 5).

Human-eagle conflicts are one of the key threats to the future survival of the Philippine Eagle (Ibañez *et al*. 2016). Due to the habitat preferences identified here for forest edges and clearings which are the same areas humans occupy, the likelihood of human-eagle encounters is high, which often results in death or severe injury for eagles. This is mainly through retaliatory trapping due to eagle predation on domestic animals, or accidental trapping in snares set by locals to capture wildlife in the forests. This is further exacerbated in forest edges because these areas are often designated as buffers or multiple use zones in protected areas which may not offer the protection needed for Philippine Eagles. Previously, conservation priorities for the Philippine Eagle have been focused on protecting nest sites in densely forested areas (Sutton *et al*. 2022). However, whilst this is still important, we show that adult eagles spend 75-80 % of space-time outside of core nesting areas in human fragmented landscapes. Thus, promoting eagle-friendly lifestyles and values within forest communities as part of area-based conservation is also necessary at nest sites located in forest edges, along with community incentives to reduce human-eagle conflict (Ibañez *et al*. 2016).

We recognise there are limitations to our inferences due to the low sample size of individual eagles tagged. However, the financing of expensive GPS telemetry devices, along with capturing adult eagles in rugged and remote tropical forest terrain is difficult. Tagging more adult eagles, including beyond Mindanao, would allow further interpretation of the results and conclusions here. We also recognise the large differences in the number of GPS fixes between adults and the subsequent potential bias in our results. However, all our sample sizes were within the range deemed suitable for estimating home range size (Bekoff & Mech 1984; Seaman *et al*. 1999) and resource selection (Northrup *et al*. 2013). The disparity between GPS location sample size is largely due to tagged adults being deliberately killed (Ibañez *et al*. 2016) or tags failing. There is little we can do about this in the context of the current study. However, accounting for these disparities in sample size, rates, and intervals using methods such as AKDE, whilst improving GPS device setting protocols, can remedy these issues for home range estimation.

The use of modern satellite tracking devices, combined with environmental data derived from satellite remote sensing has revolutionized our collective understanding of animal movement ecology and resource selection (Seidel *et al*. 2018). Building on the analyses here by incorporating movement models using either Hidden Markov models (HMMs; Langrock *et al*. 2012) or integrated Step-Selection Functions (iSSFs; Avgar *et al*. 2016), would further identify the drivers of Philippine Eagle space and resource use from latent behavioural states and movement patterns. Rather than focusing on a single ‘best’ home range estimator, we implemented a range of robust space use estimators, along with easily interpretable resource selection functions to accommodate variation in space and resource use across individual eagles to help inform conservation management. We recommend that analysts consider various statistical approaches to animal movement data to fully explore space-time and resource use, ensuring that model outputs are interpretable to conservation managers and practitioners.

## Acknowledgements

We thank all staff and volunteers from the Philippine Eagle Foundation (PEF) who conducted fieldwork over the past four decades, including local forest guards, nest wardens and indigenous co-researchers. LJS thanks The Peregrine Fund for providing a post-doctoral research grant and we thank the M.J. Murdoch Charitable Trust for funding. The PEF would like to thank local government partners across the Philippines, and the following institutions that funded and supported the field surveys and nest monitoring: Mohammed Bin Zayed Conservation Fund, Local Government of Apayao and Calanasan, Disney Conservation, Whitley Fund for Nature, Microwave Telemetry Inc, KoEko, Forest Foundation Philippines, The Peregrine Fund, Direct Aid Program - AusAID, USAID/Phil-Am Fund, USAID/Protect Wildlife, Insular Life Foundation, GIZ-Coseram, Pacific Paints (Boysen) Philippines, Energy Development Corporation, UNDP Global Environment Fund, Italy Debt Swamp/Department of Finance, US Forest Service, San Roque Power Corporation, Cornell Lab of Ornithology, Raptor Resource Project, and the Department of Environment and Natural Resources through the Biodiversity Management Bureau and its regional and local offices (DENR Regions 2, 4, 8, 9, 10, 11, 12, and 13).

## Data Accessibility Statement

Upon acceptance the data that support the findings of this study will be made openly available on the data repository *figshare*

## Supplementary Tables

**Table S1.** Comparison of candidate movement models for each adult Philippine Eagle from Ornstein-Uhlenbeck (OU) movement patterns including both isotropic and anisotropic variants using change in Akaike’s Information Criterion corrected for small sample sizes (ΔAICc). OUF = Ornstein-Uhlenbeck foraging process, IID = Independent and identically distributed data, ΔRMSPE = root mean square predictive error, DOF = effective number of degrees of freedom.

| ID | Movement process | $\Delta AICc$ | $\Delta RMSPE$ (km) | DOF |
| --- | --- | --- | --- | --- |
| 001F | OU anisotropic | 0.00 | 0.00 | 407.58 |
|  | OUF anisotropic | 2.01 | -3.58 | 408.61 |
|  | OU isotropic | 244.67 | 79.27 | 385.83 |
|  | OUF isotropic | 246.67 | 75.22 | 386.90 |
|  | IID anisotropic | 2238.31 | -164.98 | 1487.00 |
| 002F | OU anisotropic | 0.00 | 0.00 | 283.00 |
|  | OUF anisotropic | 2.01 | -5.10 | 284.11 |
|  | OU isotropic | 88.42 | 94.78 | 263.74 |
|  | OUF isotropic | 90.42 | 91.20 | 264.43 |
|  | IID anisotropic | 2734.57 | -183.76 | 1370.00 |
| 003F | OU anisotropic | 0.00 | 0.00 | 105.95 |
|  | OUF anisotropic | 0.53 | 2.63 | 114.50 |
|  | OU isotropic | 3.49 | -16.74 | 109.38 |
|  | OUF isotropic | 4.45 | -15.60 | 116.79 |
|  | IID anisotropic | 109.75 | -28.85 | 263.00 |
| 004M | OU anisotropic | 0.00 | 0.00 | 51.41 |
|  | OUF anisotropic | 2.06 | -39.40 | 52.52 |
|  | OU isotropic | 21.52 | -182.89 | 56.90 |
|  | OUF isotropic | 23.55 | -213.36 | 57.92 |
|  | IID anisotropic | 464.52 | -439.68 | 240.00 |
| 005M | OU anisotropic | 0.00 | 0.00 | 146.62 |
|  | OUF anisotropic | 1.46 | -40.04 | 148.61 |
|  | OU isotropic | 367.55 | -802.23 | 147.62 |
|  | OUF isotropic | 366.94 | -851.92 | 152.08 |
|  | IID anisotropic | 23879.18 | -2579.50 | 5274.00 |
| 006F | OU anisotropic | 0.00 | 0.00 | 59.86 |
|  | OUF anisotropic | 1.98 | -41.83 | 60.86 |
|  | OU isotropic | 160.71 | 380.38 | 51.87 |
|  | OUF isotropic | 162.68 | 329.77 | 52.83 |
|  | IID anisotropic | 14844.51 | -54.53 | 2952.00 |

## Supplementary Figures

**Figure S1.**
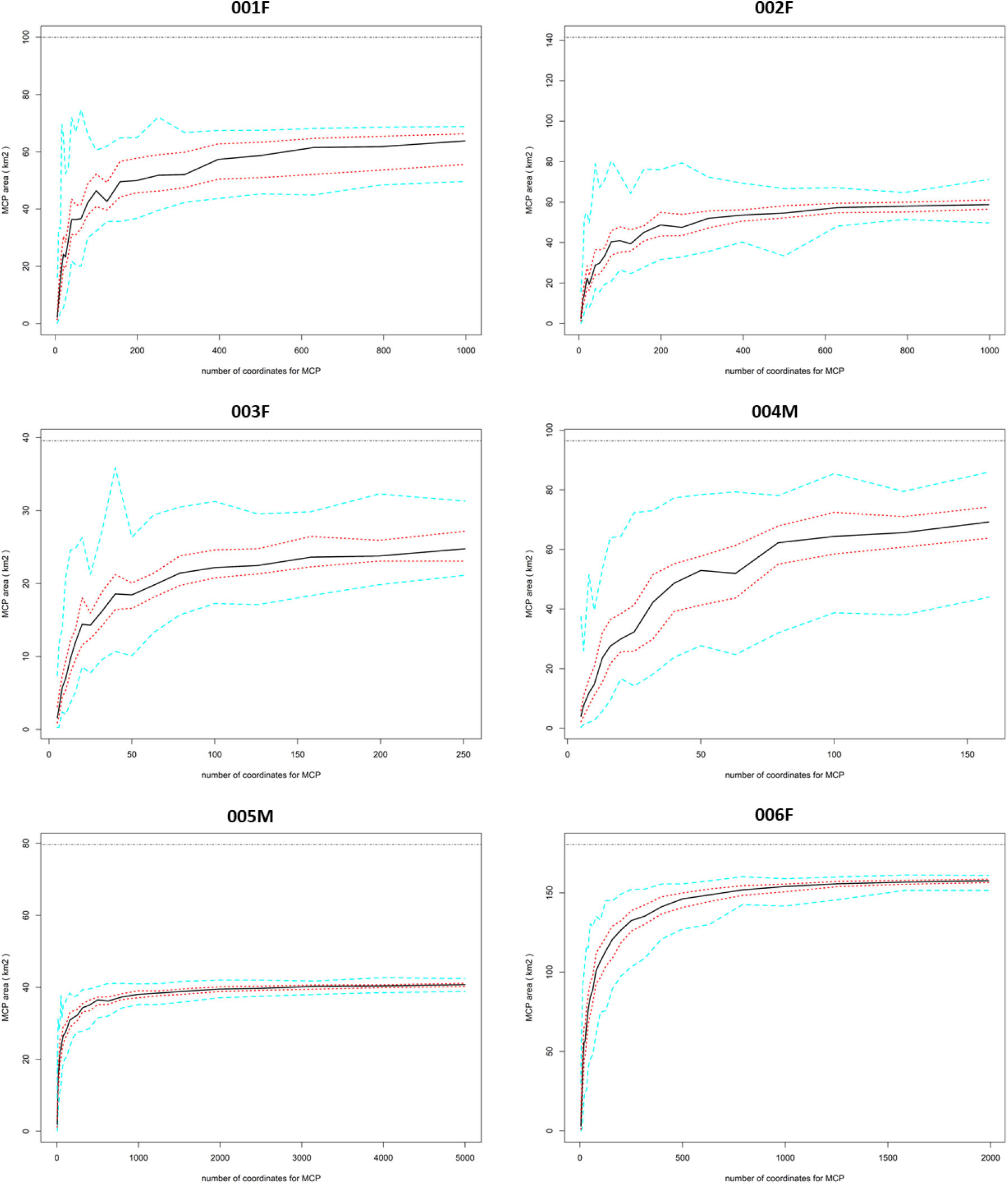
Incremental analysis using bootstrapped minimum convex polygons (*n =* 100), quantifying when the number of GPS relocations within the MCP area reached an asymptote for capturing the utilization distribution for six adult Philippine Eagles on the island of Mindanao. Black line indicates 50% percentile of MCP area, dashed red lines lower 25% percentile and upper 75 % percentile of MCP area and dashed turquoise lines indicate 0% and 100% percentile of MCP area.

**Figure S2.**
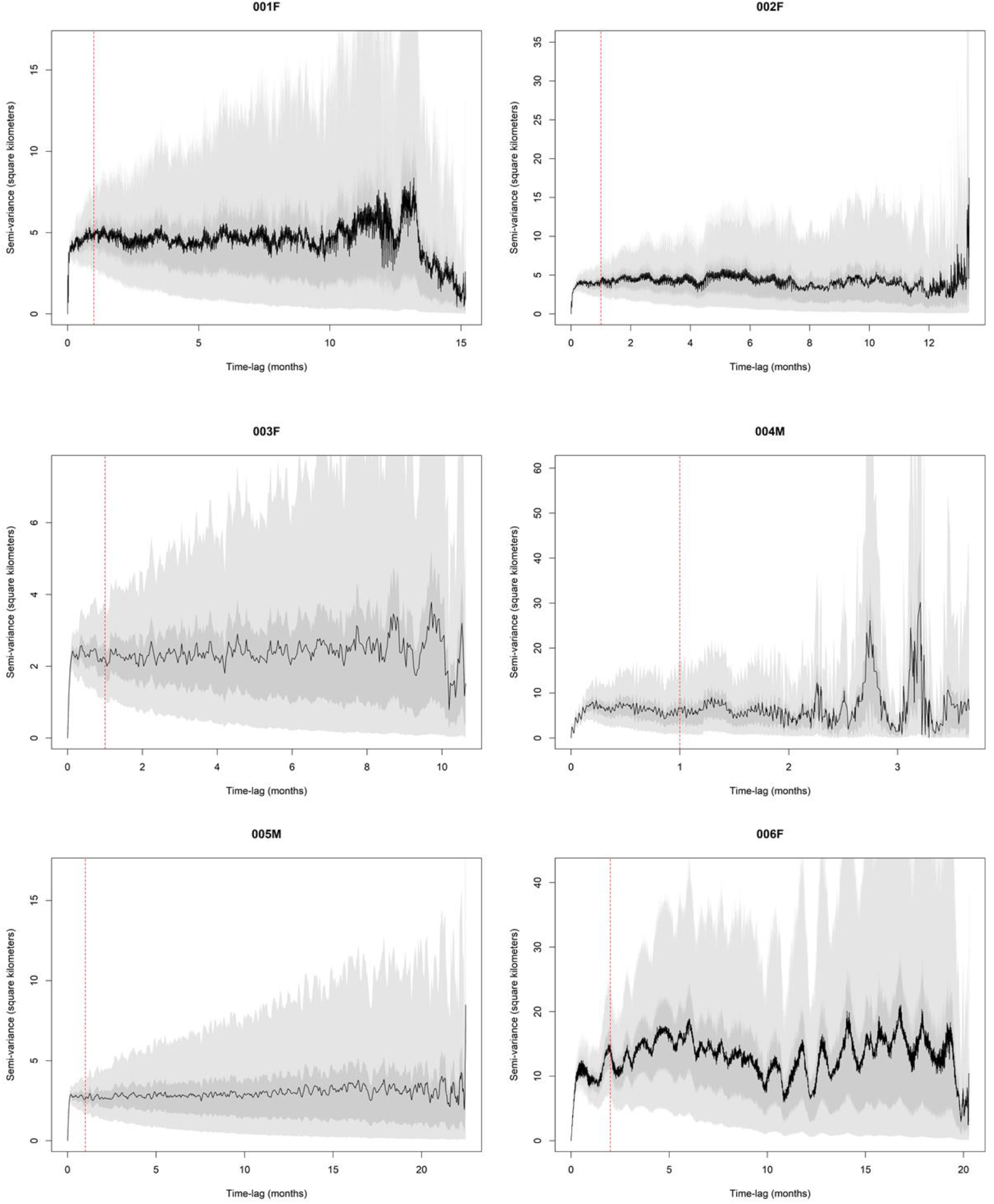
Range residency tests calculated over the entire sampling period for six adult Philippine Eagles on the island of Mindanao using semi-variance functions visualised with empirical variograms to identify unbiased estimates of stationary movement periods of site fidelity. Red vertical line indicates range residency asymptote with Markovian Confidence Intervals for calculating the maximum number of non-overlapping lags.

**Figure S3.**
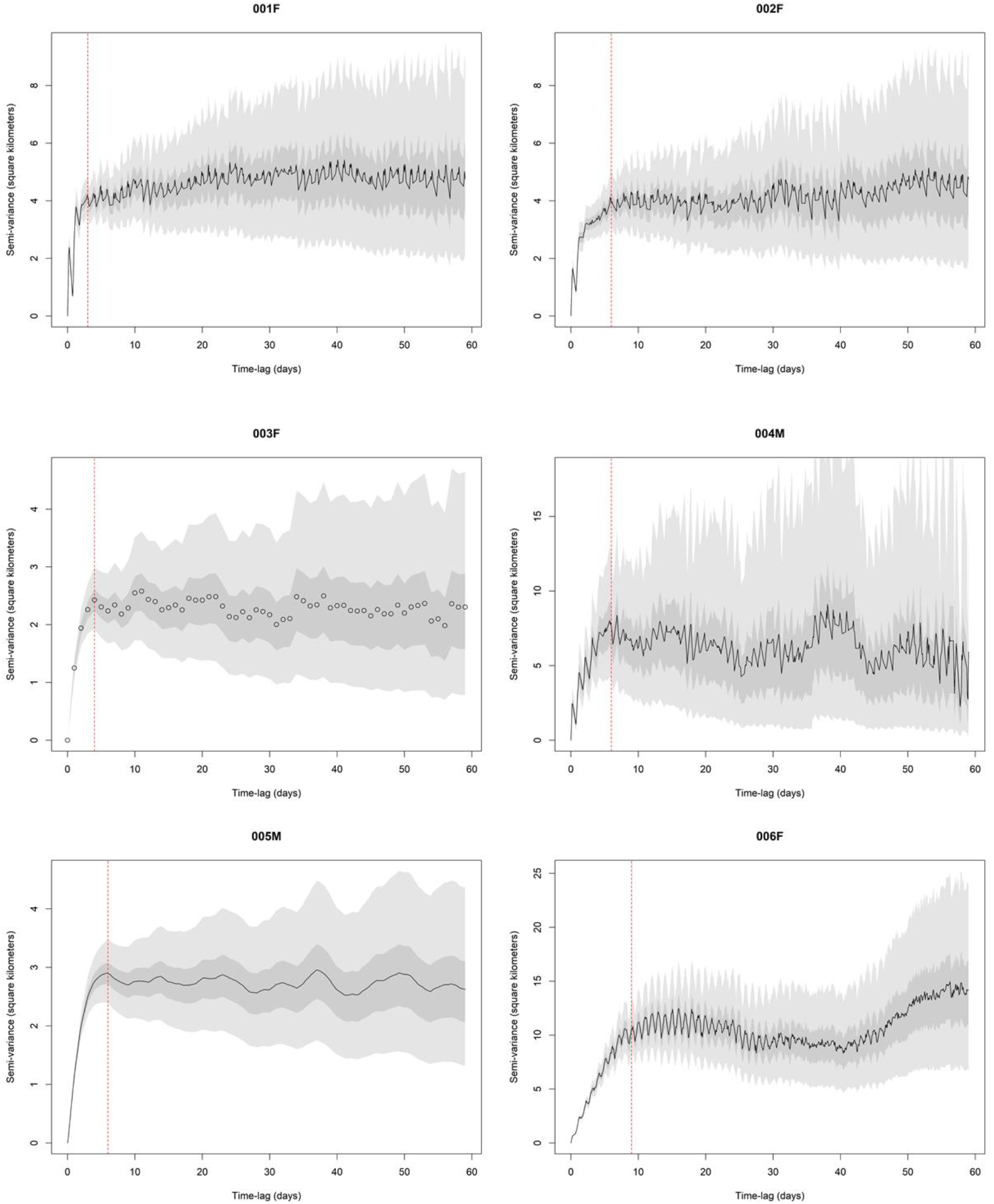
Range residency tests calculated over a 60-day sampling period for six adult Philippine Eagles on the island of Mindanao using semi-variance functions visualised with empirical variograms to identify unbiased estimates of stationary movement periods of site fidelity. Red vertical line indicates range residency asymptote with Markovian Confidence Intervals for calculating the maximum number of non-overlapping lags.

**Figure S4.**
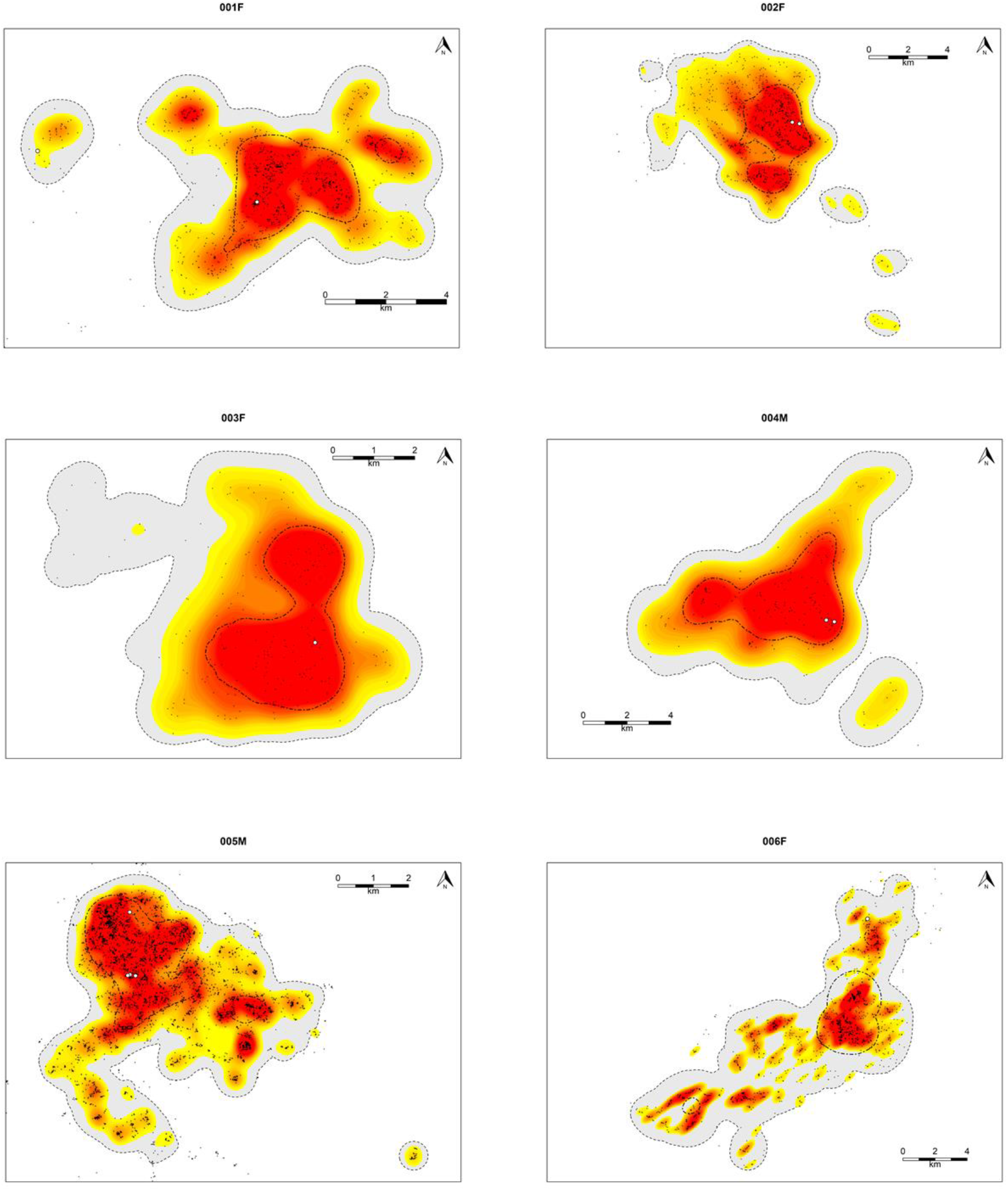
Fixed and adaptive kernel density estimates for six adult Philippine Eagles on the island of Mindanao. Fixed kernel estimates calculate 95 % probability of use (grey with hashed border) to represent the home range utilization distribution and 50 % probability of use (black dot-dash line) to represent a core range utilization distribution. Adaptive kernel estimates calculate 95 % probability of use contour isopleths (red) to represent the home range utilization distribution and 50 % probability of use contour isopleths (yellow) to represent a core range utilization distribution. Black points show filtered locations using a 3-hr sampling interval for each respective adult Philippine Eagle. White points indicate nest sites.

**Figure S5.**
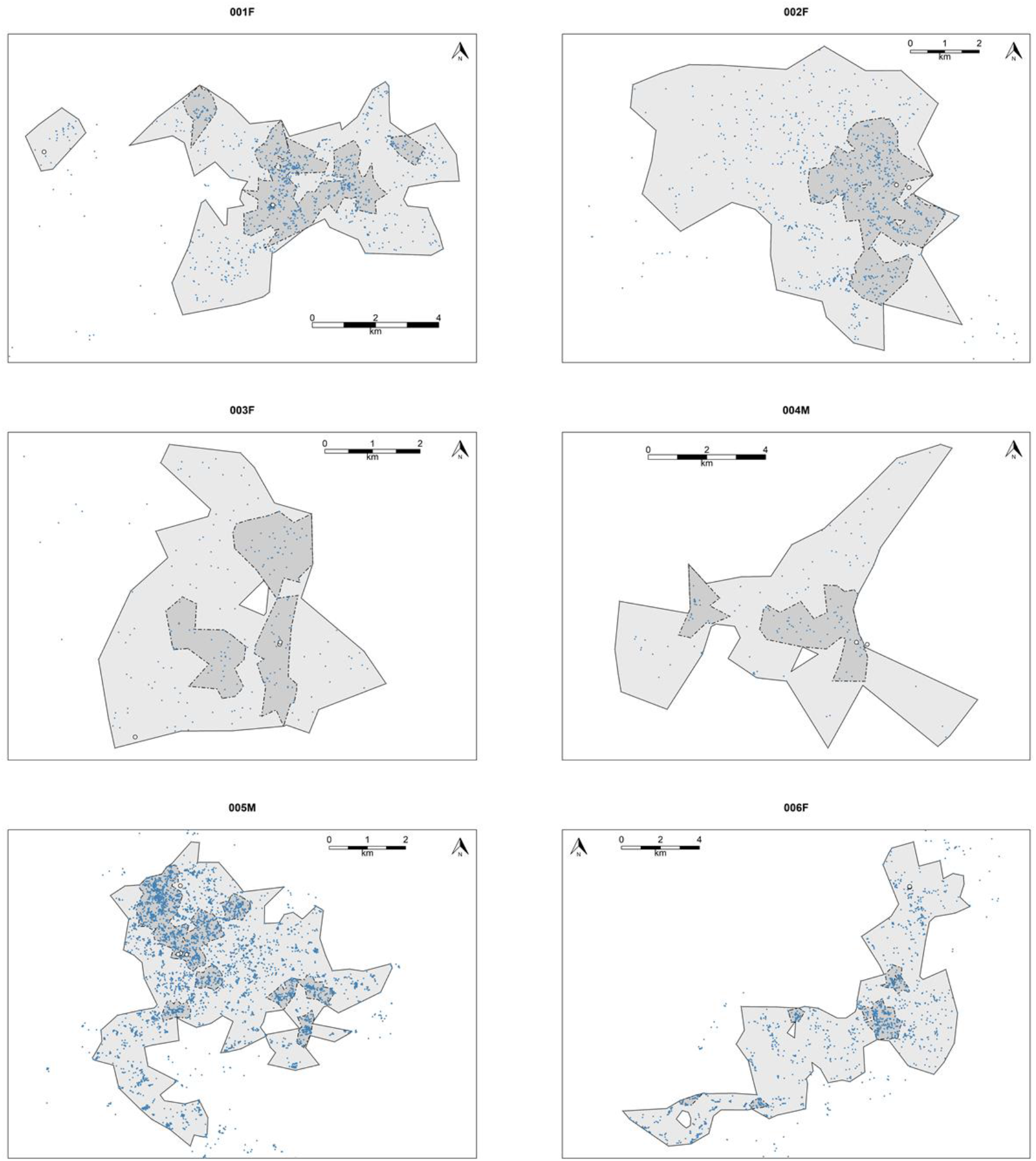
Local Convex Hull (LoCoH) home range estimates for six adult Philippine Eagles on the island of Mindanao. Estimates calculate 95 % probability of use to represent the home range utilization distribution (light grey) and 50 % probability of use to represent a core range utilization distribution (dark grey). Blue points show filtered locations using a 3-hr sampling interval for each respective adult Philippine Eagle. White points indicate nest sites.

